# The Molecular Basis for Hydrodynamic Properties of PEGylated Human Serum Albumin

**DOI:** 10.1101/2024.03.05.583576

**Authors:** Patrick J. Fleming, John J. Correia, Karen G. Fleming

## Abstract

Polyethylene glycol conjugation provides a protective modification that enhances the pharmacokinetics and solubility of proteins for therapeutic use. A knowledge of the structural ensemble of these PEGylated proteins is necessary to understand the molecular details that contribute to their hydrodynamic and colligative properties. Because of the large size and dynamic flexibility of pharmaceutically important PEGylated proteins, the determination of structure is challenging. Here we demonstrate that structural ensembles, generated by coarse-grained simulations, can be analyzed with HullRad and used to predict sedimentation coefficients and concentration dependent hydrodynamic and diffusion nonideality coefficients of PEGylated proteins. A knowledge of these properties enhances the ability to design and analyze new modified protein therapeutics.

**STATEMENT OF SIGNIFICANCE:** Proteins constitute a growing class of biotherapeutics. Chemical modification(s) with inert polymers are known to enhance the serum half-life and formulation of these biological therapeutics but the effects of modification on protein-protein interactions in solution have been difficult to predict. Here we describe methods for predicting the molecular basis for the hydrodynamic properties of polymer conjugated proteins that determine their solution behavior.

## INTRODUCTION

The conjugation of polyethylene glycol (PEG) to proteins, also called PEGylation, is a useful modification in the biopharmaceutical industry for extending serum half-life and improving formulation of protein therapeutics (1–3). Understanding how PEGylation changes the chemical and bioactivity properties of proteins requires a knowledge of the structures of these conjugates. However, investigation of the structure of PEGylated proteins is difficult because the PEG is flexible and explores a large conformational space. Although time and ensemble averaged properties may be obtained with sedimentation or scattering methods, molecular details are more difficult to determine. All-atom molecular simulations have been used to elucidate atomic configurations for relatively small PEG-protein conjugates (4–7), however, these are less useful for the large PEG molecules that may be used in protein therapeutics. Therefore, there is a need for computational methods that can bridge the gap between all-atom simulations and solution measurements.

Our goal is to obtain the calculated properties of molecular model ensembles of PEG-protein complexes as described in this report to visualize the structure, and understand the hydrodynamic and thermodynamic properties, of the same PEG-protein conjugates as measured by analytical ultra-centrifugation (AUC) and dynamic light scattering (DLS) in the companion paper (8). AUC measures the sedimentation coefficient *s*^0^, relative shape *f/f*_o_, and total effective hydration *V*_s_*/v* of macromolecules that give rise to a Stokes radius *R*_S_. In addition, AUC measures the concentration dependence of sedimentation in terms of hydrodynamic nonideality *k*_S_ and thermodynamic nonideality *BM*_1_, while DLS measures the concentration dependence of diffusion *k*_D_. Collectively, these parameters represent the impact of shape and effective hydration on the transport, diffusion, and colligative solution properties of PEG-protein complexes.

Modifications to the hydrodynamic theory implemented in the Hullrad algorithm described here allow an investigator to calculate these properties from a model ensemble of random coil polymer-protein complexes. These modifications accurately predict the experimental results while also elucidating the underlying principles that dictate the colligative properties of these therapeutic macromolecules.

We generated molecular models of several sizes of PEGs and PEGylated human serum albumin using a coarse-grained modeling protocol. The resultant structural ensembles from molecular simulation trajectories were used to investigate the calculated molecular properties that contribute to the experimental results found in the companion paper (8). Notably, a simple coarse-grained model accurately reproduces the fundamental hydrodynamic properties of PEGylated human serum albumin (PEG-HSA). The results highlight the different contributions of PEG and protein to the overall hydrodynamic properties of the conjugate. A major finding is that changes in total hydration explain the different sedimentation and diffusion properties of PEGylated proteins. The new modifications to the HullRad algorithm result in accurate calculations of macromolecular colligative properties that are important in the formulation of protein therapeutics.

## METHODS

### Coarse-Grained Model and Simulation

For short chain PEG it is possible to perform all-atom molecular simulations (4–6). But for the longer PEG chains of interest in the biopharmaceutical industry, coarse-grain (CG) simulations are necessary to obtain Boltzmann distributions of conformations within a reasonable time. A CG model for simulation of PEG alone in explicit water has been reported (9). But the PEG-HSA conjugates studied in the companion paper range in size from 5 kDa to 40 kDa PEG. These large polymers would require a prohibitively large amount of explicit water to accommodate extended conformations. Therefore, we elected to use a simulation protocol *in vacuo*.

We chose a 3 kDa PEG size (PEG68) for initial model validation because it is at the upper range of recent experimental characterization (10) and in the middle range of previous CG molecular simulations (9). Model ensembles of PEG alone and PEG-HSA were generated using CafeMol (11). This open-source molecular dynamics application uses either C_*α*_-only or C_*α*_–C_β_ coarse-graining for polypeptide and nucleic acid chains in vacuo. We included PEG in CafeMol by using a single C_*α*_ pseudo-atom CG model constructed as described below.

A linear PEG68 (68 ethylene oxide units) CG model consisting of 68 pseudo-atoms connected with bond length 3.7 Å was built in PyMOL (12). Bond length is based on the C1-C1 distance in an all-atom model of PEG built with CHARMM-GUI (13). The CG model is a polymer with the ethylene oxide units represented by single spheres centered on the C1 atom as illustrated in Fig. S1. Chain end units were treated as being identical to interior units. This linear polymer model was the starting structure for the simulations carried out during parameterization of the CafeMol excluded volume term as described below. Multiples of the PEG68 model were used to build larger PEG structures and PEG-HSA conjugates with PyMOL.

The coarse-grained model of PEG-HSA conjugate was designed to match the conjugate described in the companion paper (8) and is illustrated in Fig. 1. The number of ethylene oxide groups in each conjugate are: 114, 227, 454, 908 for 5K PEG-, 10K PEG-, 20K PEG-, and 40K PEG-HSA, respectively X-ray crystal structural models of HSA contain missing residues. For this reason, PEG-HSA conjugate models were generated using the AlphaFold2 (14) model AF-P02768-F1 of HSA. AF-P02768-F1 is an excellent match to existing X-ray crystal structures: Corresponding C_*α*_ atom RMSD with crystal structure 4F5S = 1.18 Å, with 1A06 = 0.69 Å, with 1E7B = 0.53 Å, and with 5Z0B = 0.89 Å. The signal and prepro sequences (residue numbers 1-24) were removed from the model to obtain the mature protein sequence. The PEG-protein conjugate was built using PyMOL by attaching the succinimide residue to HSA cysteine 34. The acetamide and succinimide residues were modeled as additional PEG units (cf. Fig. 1).

**Figure 1.**
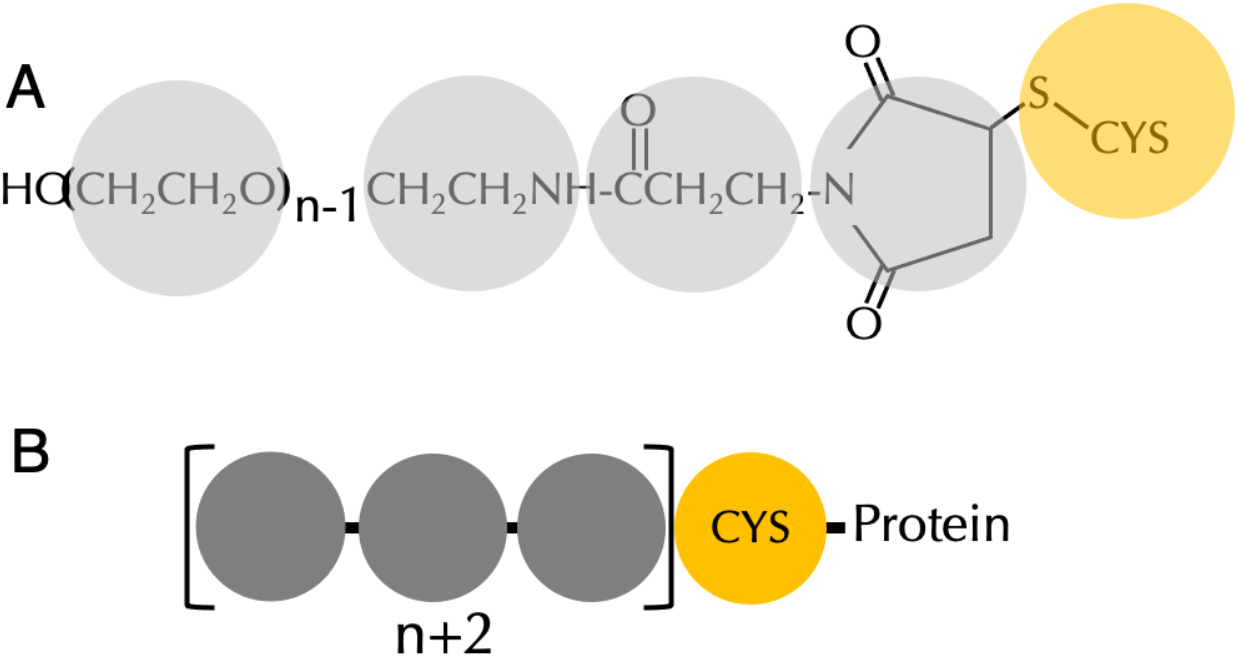
PEG-protein conjugate coarse-grained model. (A) Structure of polyethylene glycol maleimide thioether conjugated to a protein cysteine residue. Coarse-grained pseudo-atoms representing the chemical groups are shown as colored circles. The number of ethylene oxide groups in a PEG species is n. (B) Schematic diagram of CafeMol coarse-grained PEG-protein conjugate model. Colors correspond to those in panel (A).

Coarse-grained simulations of PEG and PEG-HSA conjugates were run at 293.15 K using Langevin dynamics with residue-specific mass (11), excluded volume repulsive interaction (Eq. 1), and local bond (Eq. 2) potentials,

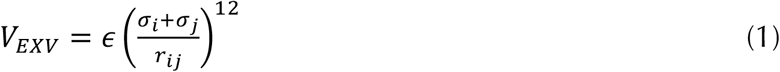

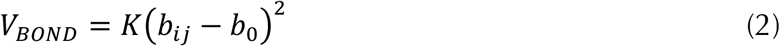

where *ε* and *K* are relative energy coefficients, σ_*i*_ is the CG pseudo-atom *i* excluded volume radius, *r*_*ij*_ is the distance between pseudo-atoms *i* and *j, b*_*ij*_ is the instantaneous bond length between pseudo-atoms *i* and *j*, and *b*_*0*_ is the ideal bond length. CafeMol default values were used for all parameters except for the PEG pseudo-atom σ and *b*_*0*_ as described below. We emphasize that there is no attractive interaction potential in the force field used here and that the dynamics of the PEG are unrestrained. This is in contrast to our previously reported CafeMol simulations of unfolded polypeptides (15).

The protein portion of the conjugate is converted to single pseudo-atom residues centered on the C_*α*_ atom by CafeMol for simulation. Masses, bond lengths and excluded volumes for CG amino acid residues were the default values in CafeMol. For PEG the mass was set at 44.05 g/mol per unit; bond length was 3.7 Å, and an excluded volume radius of 3.08 Å was empirically determined as described below. Step size was 0.4, and total simulation steps were 2.5 x 10^7^ to 1.0 x 10^8^ depending on the size of the PEG alone or conjugate as described below. Fig. S4 shows the time evolution of calculated *R*_*G*_ during simulations of PEG68 and the largest PEG studied, PEG908 (40K PEG). We found that 2.5 x 10^7^ steps of simulation were sufficient for convergence of the PEG68 system properties as expected from previous CafeMol simulations of unfolded polypeptides (15). As shown in panel (B) of Fig. S4 the much larger 40K PEG requires a longer simulation time to reach equilibrium. A simulation time of 1.0 x 10^8^ steps was run for 40K PEG and 40K PEG-HSA and a simulation time of 5.0 x 10^7^ steps was found to be adequate for convergence of 5K PEG-, 10K PEG-, and 20K PEG-HSA, respectively.

Three independent simulations were run for each molecular species. The first 20% of trajectories were considered equilibration and 1000 frames from the remaining trajectory were evenly sampled to generate ensembles for analysis. Identical results were obtained for ensembles of 1000 and 2000 frames for the largest PEG (40K PEG) and therefore ensembles of 1000 structures were analyzed for all species. For some analyses reported below the three ensembles were combined. Previous studies on PEGylated lysozyme showed no effect of the conjugated PEG on the structure of the protein (16), therefore, for the PEG-HSA simulations the protein residues and the pseudo-atom representing succinimide were fixed in position, and only the PEG and acetamide were unrestrained. This essentially created a bond between the PEG succinimide end and the protein. Model ensembles of various sizes were output from the trajectories by sampling at intervals using CATDCD and VMD (17).

### Calculation of Hydrodynamic Properties

The fundamental calculation of HullRad is the hydrodynamic volume of a molecular model using a convex hull (18). This volume includes the molecular atomic volume, first shell hydration volume, and entrained water volume (19). The product of the radius of an equivalent hydrodynamic volume sphere and a Perrin-like shape factor gives the Stokes radius, *R*_*S*_. From the *R*_*S*_, HullRad calculates diffusion coefficients 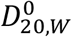 from the Stokes-Einstein-Sutherland equation (Eq. 3) and sedimentation coefficients 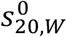 from the Svedberg equation (Eq. 4),

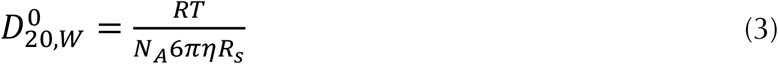

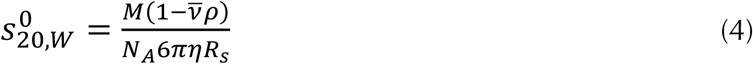

where the subscript 20,W indicates 20°C in water, the zero superscript indicates that these are properties at infinite dilution, *R* is the gas constant, *T* is temperature, *N*_*A*_ is Avogadro’s number, *η* is the solvent viscosity, *M* is molecular mass, 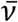 is the molecular partial specific volume (not including hydration water), and *ρ* is the solvent density. In the remaining text the zero superscript is omitted for convenience.

HullRad was extended to calculate hydrodynamic properties of PEG directly from the coarse-grained model using a partial specific volume of 0.83 (20). PEG-HSA models were analyzed after the protein portion of the CG model was substituted with the AF-P02768-F1 all-atom model for analysis with HullRad. Superposition of the all-atom protein model on the CG simulation model was accomplished with VMD. Axial ratio *a/b* is calculated by HullRad from an ellipsoid of revolution fit to the convex hull volume and shape.

## RESULTS and DISCUSSION

### Parameterization of Coarse-Grained Model

The only adjustable parameters in the CG model used here are the bond length and CG pseudo-atom radius (i.e., excluded volume σ). Bond length between the PEG CG pseudo-atoms is set to be equivalent to the distance between ethylene oxide C1 atoms in an all-atom model (13). To calibrate the PEG excluded volume σ we ran simulations with varied σ values and compared the ensemble average calculated sedimentation coefficients to experimentally reported sedimentation coefficients of PEG (21,22) as shown in Figs. S2 and S3. An excluded volume σ of 3.08 Å for the PEG residue was used in all simulations.

### PEG Molecular Models are Random Coils and Agree with Experimental Radius of Gyration

For a random coil, the end-to-end distance distribution (*D*_*ee*_) is Gaussian and the relationship of *R*_*G*_ to *D*_*ee*_ is described by Eq. 5 (10).

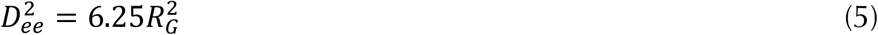

Fig. 2 shows a Gaussian distribution of *D*_*ee*_ for a combined PEG68 ensemble. Using Eq. 5 the calculated *R*_*G*_ is 1.80 nm

**Figure 2.**
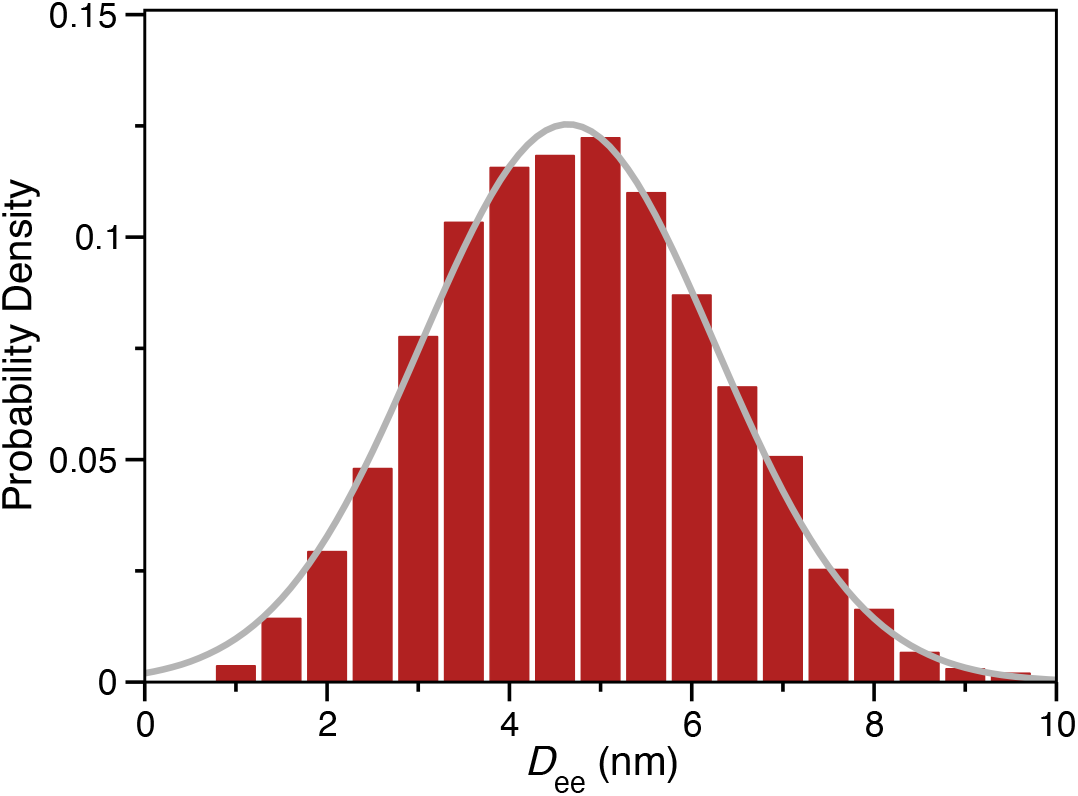
PEG68 End-to-end distance describes a random coil in solution. The distribution of end-to-end distances for the combined ensemble of PEG68 models is plotted as a histogram (*N*=3000). The grey line is the best fit to a Gaussian distribution.

The anhydrous radius of gyration (*R*_*G*_) may also be calculated using Eq. 6,

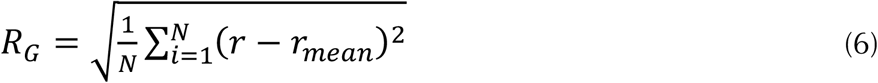

where *N* is the number of atoms (or, in the case of PEG, CG pseudo-atoms) and ***r*** is the atomic position. The combined ensemble average of 1.83 nm using Eq. 6 is consistent with the the *R*_*G*_ calculated from the distribution of *D*_*ee*_ and this agreement is evidence that the simulation ensemble is correctly modelling PEG as a random coil in solution.

The *R*_*G*_ for PEG77 has been reported by two groups using neutron scattering (23,24). We built a PEG77 CG model and generated ensembles as described above for PEG68 to further validate the model. The CG ensemble calculated *R*_*G*_ of PEG77 (using Eq. 6) agrees with experimental results obtained with neutron scattering as shown in Table S1.

As described in METHODS, HullRad uses a convex hull to calculate the hydrodynamic volume of a macromolecule. An initial convex hull is generated with the atomic centers of the molecular model. This initial hull is expanded to account for the first shell of hydration water (18). The expansion of the initial hull in HullRad was parameterized for proteins and nucleic acids. The fact that CG models of PEG resulted in calculated ensemble average values of *D*_*ee*_, *R*_*G*_, and *s*_20,W_ that are consistent with each other and agree with experimental values is evidence that the first shell hydration correction applied by HullRad is appropriate for PEG.

### PEGylated Human Serum Albumin Exhibits Large Conformational Flexibility

Fig. 3 shows a composite image of multiple frames from 5K PEG-HSA and 40K PEG-HSA simulations. The PEG moiety explores a large conformational space. Although the PEG groups do occasionally contact the protein surface as described by others (25–28), contact is transient and collisional as expected from the absence of attractive forces in the CG force field.

**Figure 3.**
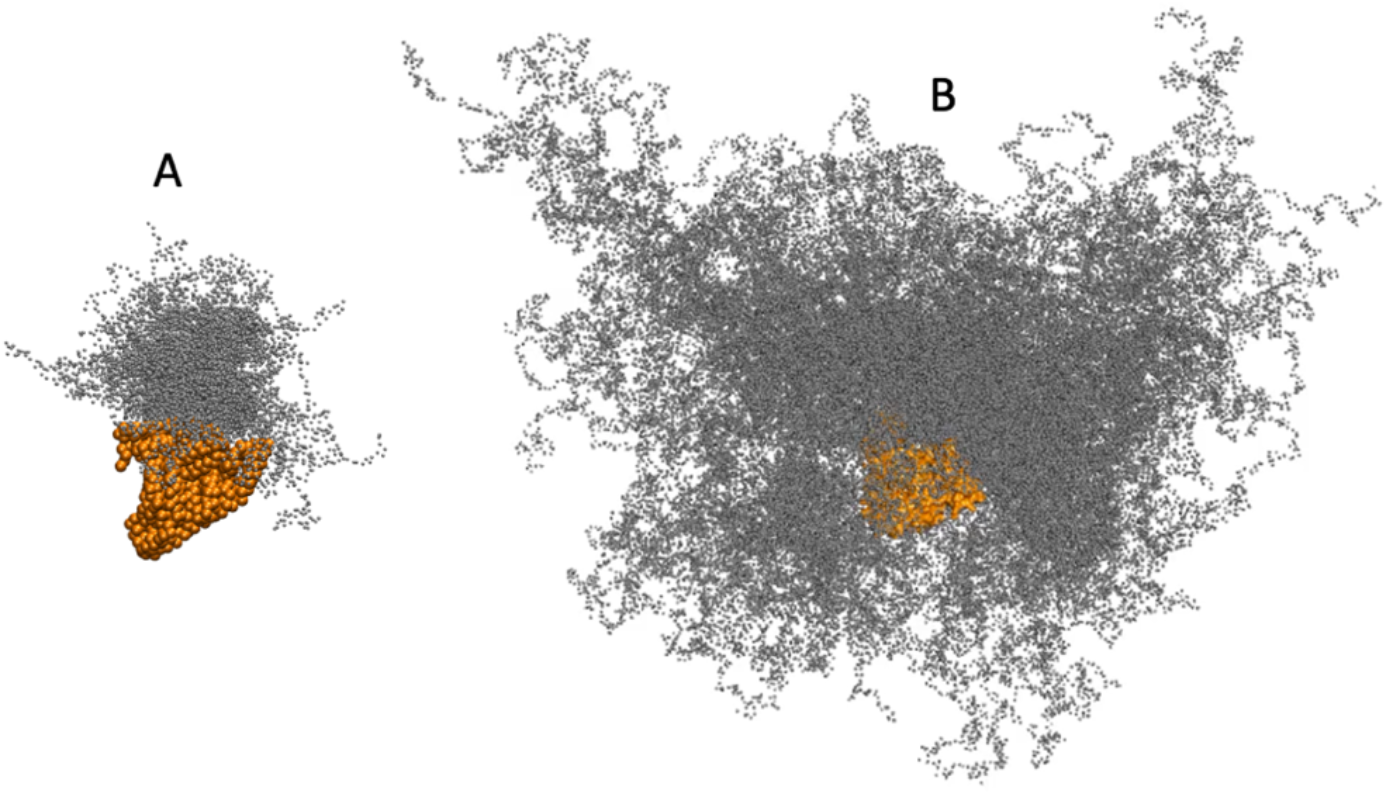
The PEG moiety of PEG-HSA samples diverse conformations. Collections of 100 evenly spaced frames from (A) 5K PEG-HSA and (B) 40K PEG-HSA simulations are overlayed in single composite images and demonstrate the extensive conformational sampling of the PEG (small grey spheres) attached to HSA (orange spheres). VMD (17) was used to create the image.

The ensemble average calculated size of the 5K PEG moiety illustrated in Fig. 3 is consistent with the experimental results reported for 5K PEG conjugated to galectin-2, a 14.5 kDa protein (29). Using small angle X-ray and neutron scattering of PEGylated galectin-2, He et al. report a 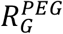 of 2.5 nm and we calculate an ensemble average 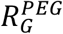 of 2.6 nm for the PEG moiety of 5K PEG-HSA.

### PEG Molecular Models Predict Fundamental Hydrodynamic Properties

The simulated ensembles described here accurately model the hydrodynamic properties of PEG-HSA conjugates at infinite dilution. The PEG and PEG-HSA hydrodynamic properties, *R*_*S*_, *s*_20,W_, and *D*_*20*,*W*_ calculated from model ensembles are compared to experimental data in Fig. 4 and detailed in Table S2.

**Figure 4.**
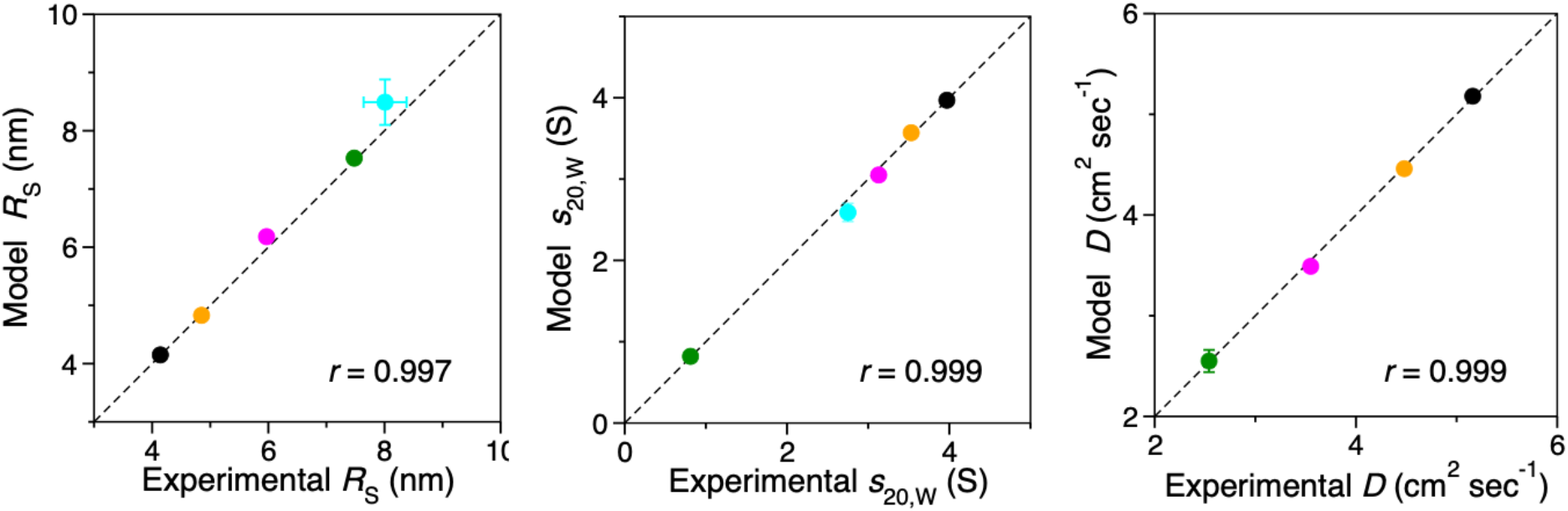
Calculated PEG and PEG-HSA hydrodynamic properties agree with experimental values. The data in Table S2 are plotted as circles. Black, 5K PEG-HSA; orange, 10k PEG-HSA; magenta, 20k PEG-HSA; green, 40K PEG-HSA; cyan, 40k PEG. Standard deviations are shown as capped error bars; some error bars are smaller than the corresponding data circle, linear regression correlation coefficients are labeled as *r*. The dashed lines are for comparison and have slopes of 1.0 and intercepts of zero.

The experimental hydrodynamic properties plotted in Fig. 4 were measured in phosphate buffered saline. Our CG model is parameterized with properties of PEG experimentally determined in water (21,22). This agreement between model and experiment is evidence that NaCl at 150 mM does not significantly affect the size of PEG in solution. Although salt may theoretically screen the dipole-dipole type interactions between PEG ethylene oxide units and affect its solution properties (30), experimental measurement of the effects of NaCl and KCl on PEG intrinsic viscosity indicate that no significant effect is observed below molar concentrations of salt (31).

The diverse PEG conformations that extend from the protein surface and illustrated in Fig. 3 result in a wide distribution of individual calculated sedimentation coefficients. Examples of specific conformations of the PEG on 5K PEG-HSA, together with their relative positions in the distribution of *s*_*20*,*W*_ values, are shown in Fig. 5.

**Figure 5.**
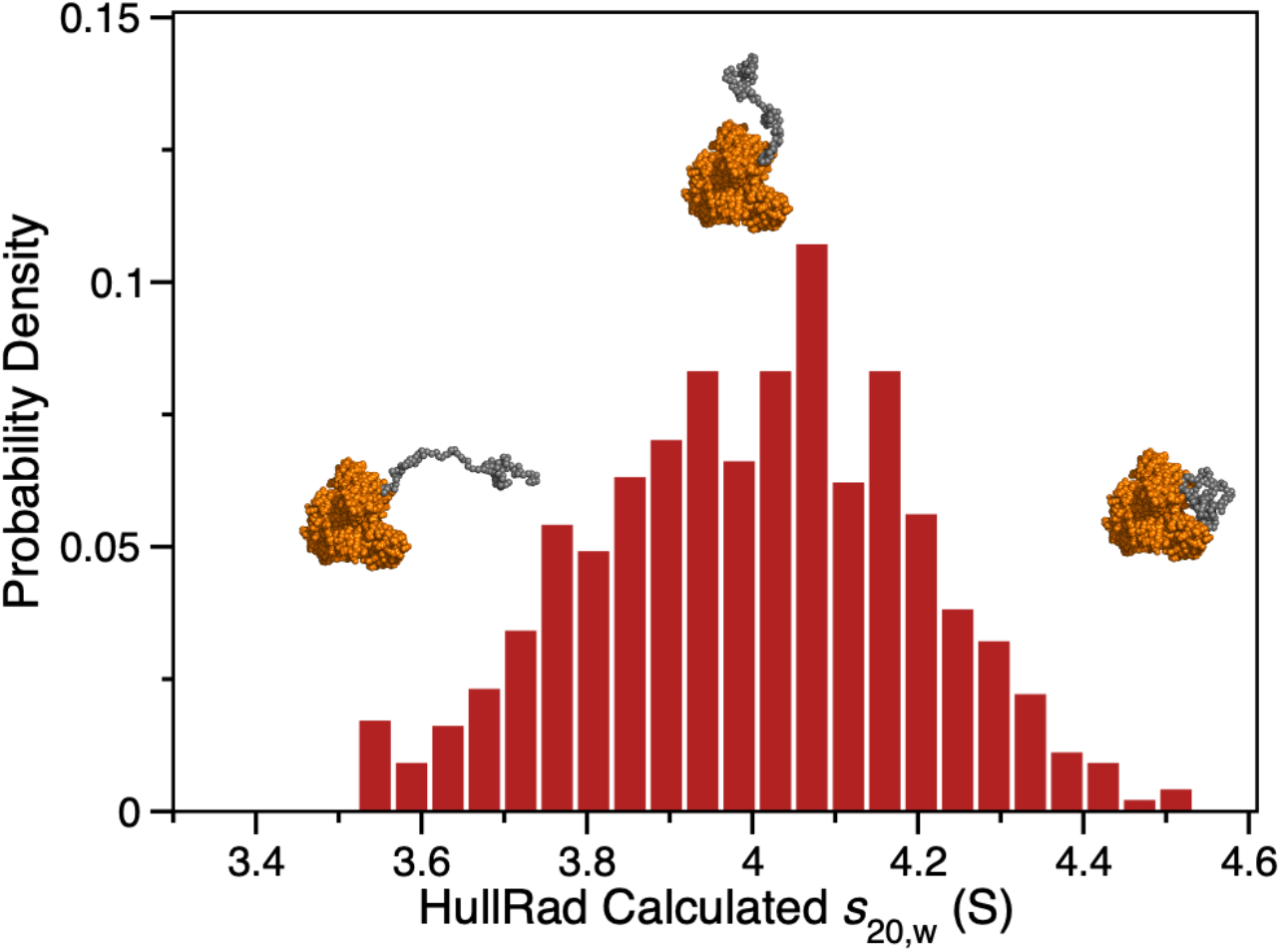
5K PEG-HSA model calculated sedimentation coefficients are widely distributed. The distribution of s_20,w_ for an ensemble (*N*=1000) of 5K PEG-HSA models is shown as a histogram plot. Several conformations of the PEGylated HSA are shown as atomic spheres with grey PEG and orange HSA; they represent conformations with the smallest, median, and largest sedimentation coefficients, respectively.

### Hydration and Shape Determine Hydrodynamic Properties

The amount of hydration water associated with a macromolecule is key to accurate calculation of the hydrodynamic properties and to the solution non-ideality discussed below (32). The hydrodynamic properties of macromolecules such as sedimentation and diffusion are determined by the effective frictional drag contributed by both hydrated molecular volume and shape. HullRad calculates the volume of a hydrated molecule using a convex hull construct. The total hydration volume is that volume within the convex hull minus the anhydrous atomic volume of the molecule and is composed of first hydration shell water and ‘entrained’ water (19). The images in Fig. 6 illustrate some examples of the initial convex hull for several conformations of a 40K PEG model (A-C) and a single PEG-HSA model conformation (D-F). For example, Figure 6C shows a large increase in entrained water (enclosed within the convex hull) as compared to Figure 6A. The HullRad calculated hydration water is much greater than that usually assumed in estimating size and shape of macromolecules from sedimentation studies interpreted with SEDNTERP (33) which effectively reports the amount of first hydration shell water. Historically, entrained water had been imagined using the term “swollen volume” (34); HullRad provides a mechanism to calculate this from structure. Two measures of hydration, the standard g/g (water/macromolecule), and the “swollen” volume (total ml/g) are listed for the molecular species in this study in Table 2. For comparison, the amount of hydration water calculated by SEDNTERP based on amino acid composition is included in the last column.

**Table 2.** Hydration of PEG, HSA and PEG-HSA models.

| Sample | HullRad<sup>a</sup> |  | SEDNTERP<sup>b</sup> |
| --- | --- | --- | --- |
| | Hydration<br>(g/g) | $V_s$<br>(ml/g) | Hydration<br>(g/g) |
| 40K PEG | 23.2 | 24.2 | 1.22 |
| 40K PEG-HSA | 12.2 | 13.1 | 0.733 |
| 20K PEG-HSA | 5.42 | 6.21 | 0.619 |
| 10K PEG-HSA | 2.69 | 3.45 | 0.540 |
| 5K PEG-HSA | 1.61 | 2.36 | 0.492 |
| HSA | 0.87 | 1.61 | 0.437 |

<sup>a</sup>Means of combined model ensembles calculated with HullRad.
<sup>b</sup>Calculated from SEDNTERP (33) or the data of Tirosh (35) as described in (8).

**Figure 6.**
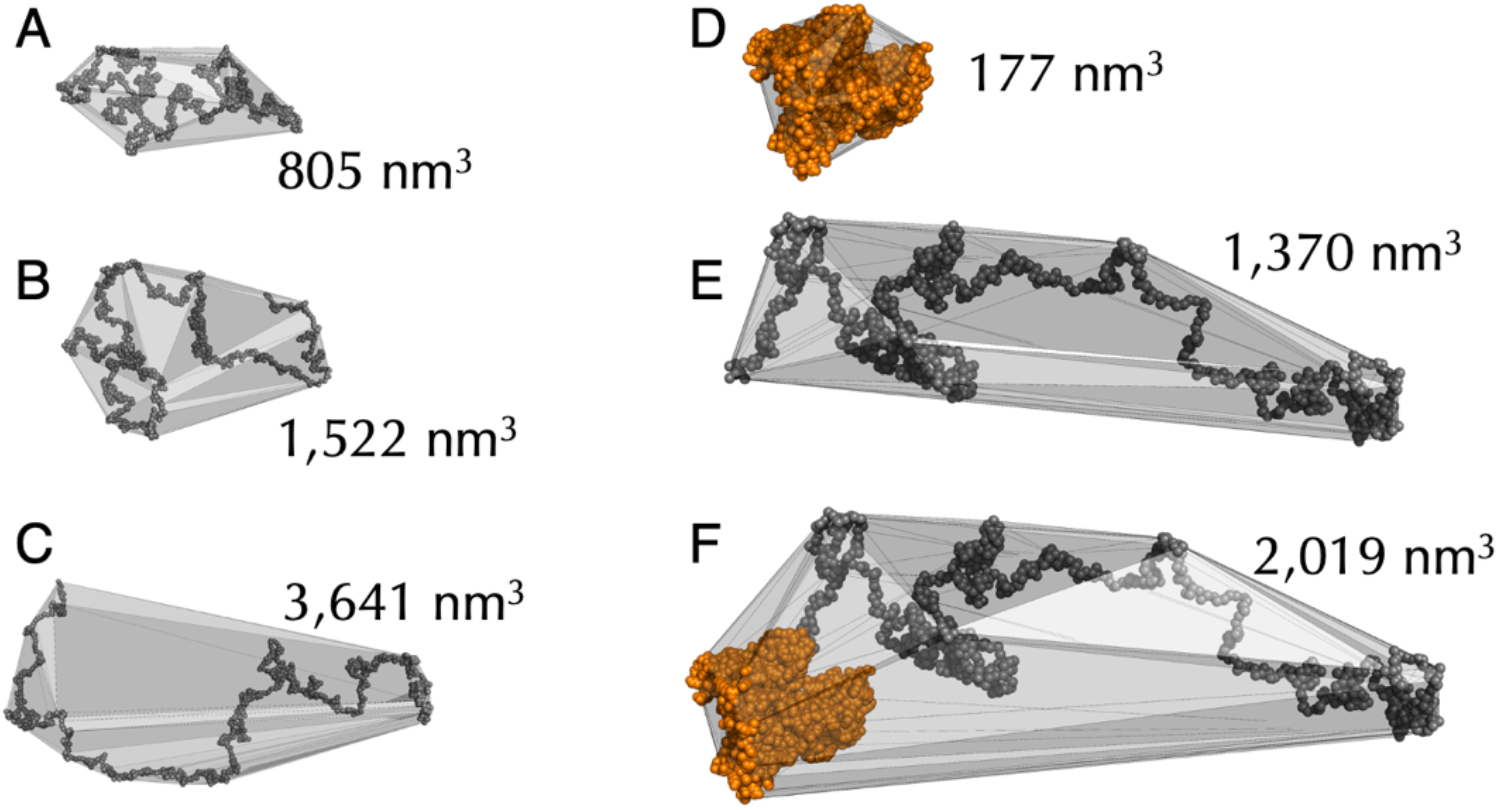
PEG and PEG-HSA hydration depends on conformation. (A) Three 40K PEG ensemble conformations with initial convex hulls and labeled with total hydration volumes. (B) A single 40K PEG-HSA species with *s*_20,w_ similar to the ensemble average (2.6 S) and showing initial convex hulls for the (Top) HSA only, (Middle) PEG only, and (Bottom) PEG-HSA conjugate; labels are for total hydration volumes. PyMOL (12) was used to create the image.

The hydration of PEG-HSA conjugates is not simply a sum of the separate PEG and protein hydration amounts. As shown in the right panel of Figure 6, additional volume is encapsulated in the convex hull of a conjugate (F) compared to a sum of the PEG moiety (E) and protein (D) convex hulls. The fact that the total hydration of the PEGylated proteins is not a sum of the separate PEG and protein hydration amounts gives rise to non-random coil scaling laws for the hydrodynamic properties of PEG-HSA conjugates. Historically, scaling laws have be used to determine whether a polymer is in a poor, neutral, or good solvent. Flory showed that for a polymer in solution, a property such as *R*_*G*_ or *R*_*S*_ follows the scaling law *R≅bN*^*v*^ where *N* is the number of residues or monomers, *b* is a constant related to persistence length, and *v* is a factor that depends on solvent quality (36). Values of *v* range from 0.33 for a collapsed polymer in a poor solvent, through 0.5 for a neutral solvent, to 0.6 in a good solvent that completely “solvates” and expands the polymer.

Fig. S5 shows log-log plots of *R*_*S*_ against molecular weight for both the PEG-HSA conjugates and the corresponding PEG moiety alone. The PEG alone plots are linear with a scaling exponent of 0.58 consistent with a random coil in good solvent. However, the corresponding PEG-HSA conjugates have an equivalent Flory scaling exponent of 1.78 indicating that their hydrodynamic size increases in a complex way with increasing molecular weight. This calculated value of *v* is consistent with the experimentally determined value of 1.63 reported in the companion paper (8). These scaling law exponents for the conjugates indicate that the increase in Stokes radius with polymer size is not a simple average of random coil and compact globular polymers.

PEG-HSA conjugates also show an unusual scaling of the radius of gyration to Stokes radius. This relationship has been of interest to facilitate the calculation of *R*_*S*_ (and therefrom, sedimentation coefficients) from experimentally determined *R*_*G*_. The relationship of *R*_*G*_*/R*_*S*_ vs *R*_*G*_ for unfolded polypeptides has been studied experimentally by Choy et al. (37) and using structural modeling by Nygaard et al. (38). The latter study showed that for a chain length less than 450 residues the relationship of *R*_*G*_*/R*_*S*_ to *R*_*G*_ is size dependent (38). The *R*_*G*_*/R*_*S*_ of the PEG-only moiety for each of the PEG-HSA conjugates is plotted against the corresponding *R*_*G*_ in Fig. S5A. These results are consistent with those of Nygaard et al. The 20K PEG (454 residues) studied here has a slope of 0.11 compared to an unfolded polypeptide of 450 residues with a slope of ∼0.1 (38). However, a similar plot for the PEG-HSA conjugates is drastically different as shown in Fig. S5B. The smallest conjugate 5K PEG-HSA has a negative slope, the 10K PEG-HSA an almost flat slope, and the larger conjugates are similar to the PEG-only plots but with smaller slopes.

The above results emphasize the unusual scaling of PEG-HSA properties with molecular size and Fig. 6D-F suggests that the complexity is related to hydration. The relationships of hydration, measured by *V*_*S*_, to sedimentation coefficients of 40K PEG and 40K PEG-HSA conjugate are shown in Fig. 7A. For the PEG-HSA conjugate (green data), the sedimentation coefficient approaches that of HSA alone (*s*_*20*,*W*_ = 3.97 S) as the *V*_*S*_ decreases and the conjugate becomes more collapsed. As the *V*_*S*_ increases, the PEG dominates the sedimentation rate (compare to cyan data). In addition, the sedimentation coefficient is affected by shape. Fig. 7B shows the combined influence of the axial ratio *a/b* and *V*_*S*_ (compare color bar) on 40K PEG-HSA *s*_*20*,*W*_.

**Figure 7.**
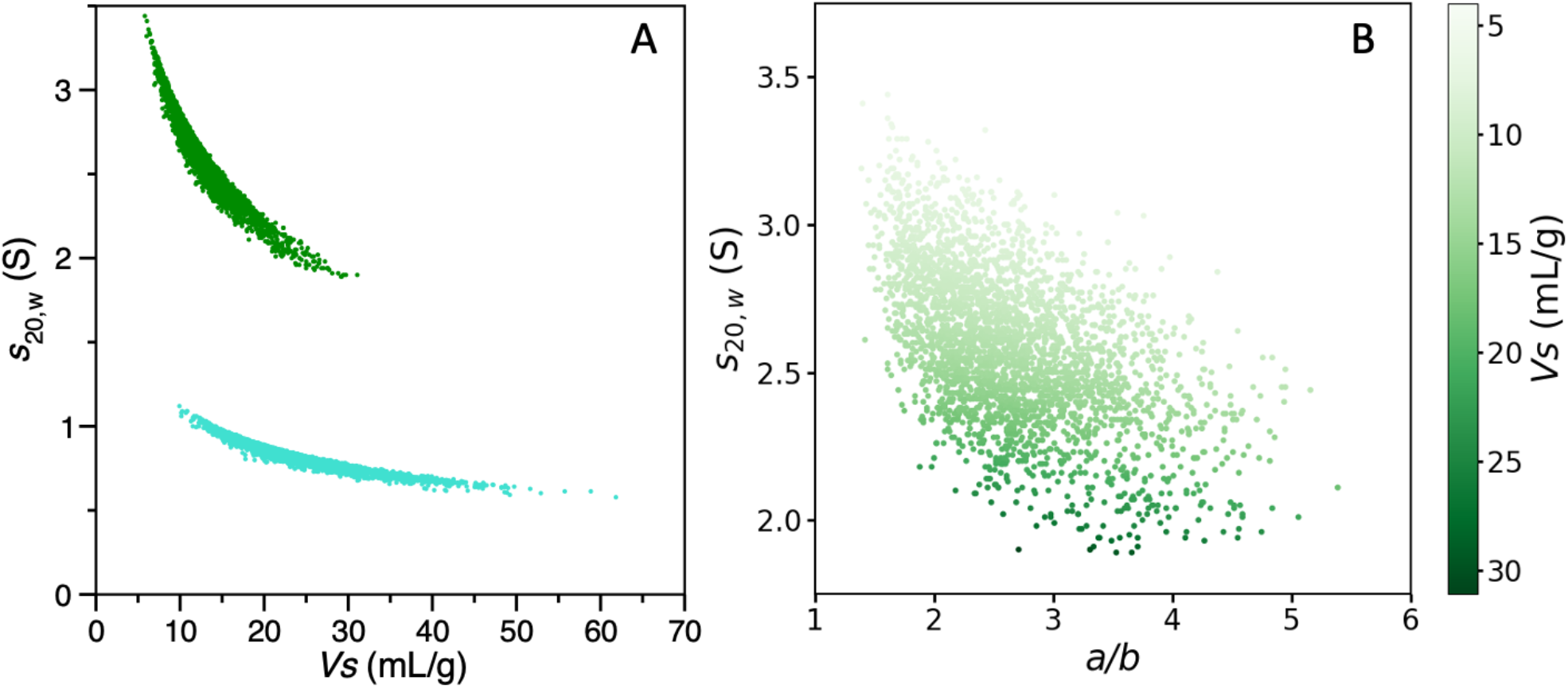
Non-linear dependence of sedimentation coefficient on hydrated volume and shape. (A) The individual model calculated *s*_*20*,*W*_ values are plotted against the corresponding hydrated (swollen) volumes *V*_*S*_ for combined ensembles (N=3000) of 40K PEG-HSA conjugate (green) and 40K PEG (cyan) and as filled circles. For comparison, the same plot for other conjugates is in Fig. S6. (B) The 40K PEG-HSA *s*_*20*,*W*_ data are plotted against the axial ratio (*a/b*). The color gradient indicates the corresponding *V*_*S*_ for each 40K PEG-HSA model in the ensemble.

The combined effects of hydration and shape on the solution properties of PEG have been investigated for many years. But historically it was not possible to independently determine both hydration and shape and this was called the “Hydration Problem” by Harding (39). For example, Kim et al. concluded that the solvent excluded volume of PEG was best calculated by a rod-like model (40) based on studies of PEG intrinsic viscosity carried out by Thomas and Charlesby (41). The hydration of PEG for the study by Kim et al. was estimated by measuring the non-freezable bound water using differential scanning calorimetry. The subsequent calculated volume fractions required the inclusion of a large shape factor to agree with experimental excluded volume measurements. However, the freezable bound water includes only the first shell of hydration water and not the entrained water that is necessary to describe the complete hydrated volume of a molecule like PEG (19).

Both Kim et al. and Thomas and Charlesby concluded that PEG deviates from a sphere and that the axial ratio increases with molecular weight to account for the increased excluded volume relationships. However, an experimental study subsequent to that by Kim et al. on the viscosity of various molecular weight PEG solutions has shown PEGs to be random coils (42) in agreement with the more recent results from sedimentation (16) and double electron−electron resonance spectroscopy (10).

In agreement with the model of PEG as a random coil, the calculated ensemble average axial ratios for the PEG-only moieties studied here do not change with PEG size, only the swollen volume correlates with increased hydrodynamic size (Fig. 8). Our combined results demonstrate that the increase in hydration with polymer size can fully explain the solution properties of increasingly large PEG without changing the axial ratios.

**Figure 8.**
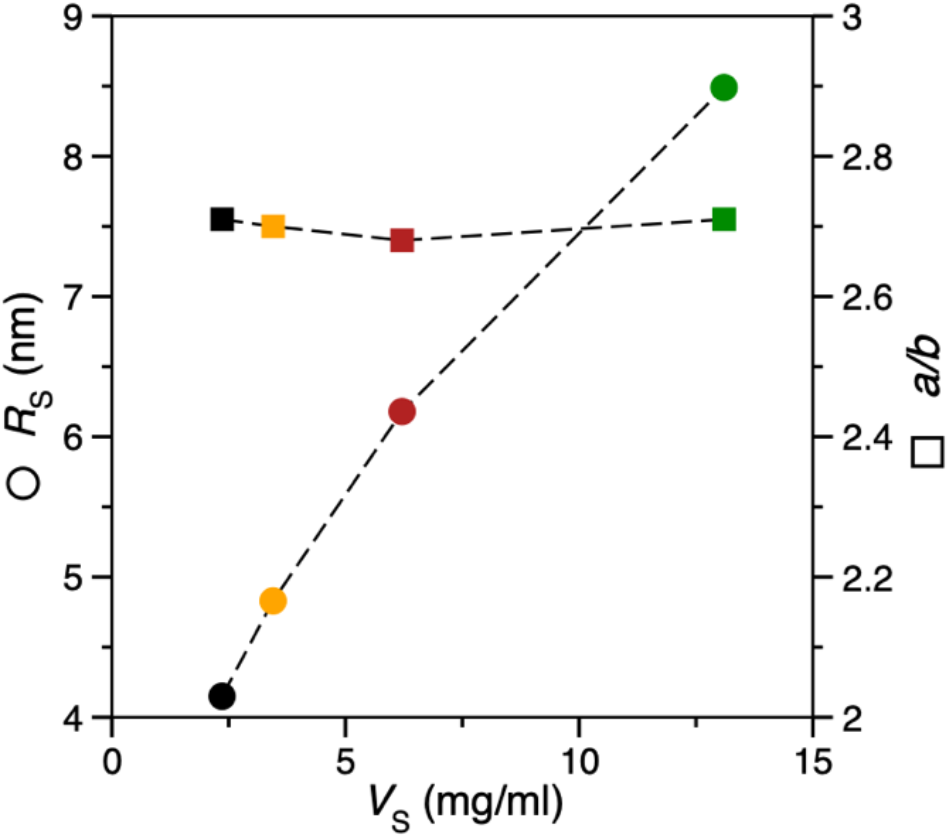
The increased Stokes radius of large PEG-HSA is determined by swollen volume increase, not shape changes. The Stokes radii *R*_*S*_ (left axis, circles) of PEG-HSA conjugates and axial ratio *a/b* (right axis, squares) of the corresponding PEG moieties are plotted against the swollen volume *V*_*S*_ of the PEG-HSA conjugates. Black, 5K PEG-HSA; orange, 10K PEG-HSA; red, 20K PEG-HSA; green, 40K PEG-HSA.

### Solution Non-Ideality

Predicting the properties of protein solutions at high concentrations is important for understanding crystallization (43), biotherapeutic formulation (44), and cellular liquid-liquid phase transitions (45). A useful approach to obtain information on the state of concentrated solutions is to determine the second virial coefficient *B*_*2*_ at semi-dilute concentrations. The second virial coefficient can be thought of as a measure of solution non-ideality. Expressions for the description of non-ideality frequently follow the form of the osmotic pressure virial equation of state (46),

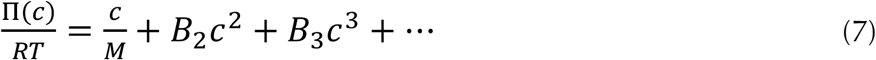

where Π is the osmotic pressure, *R* is the ideal gas constant, *T* is the temperature, *c* is the molecular concentration, *M* is the molecular mass, *B*_*2*_ is the second virial coefficient, and *B*_*3*_ is the third virial coefficient. In practice the series is truncated after the second virial term and higher order interactions are ignored except at very high concentrations. In the analytical ultracentrifugation community, the second virial coefficient is often multiplied by the molecular mass and is referred to as *BM*_*1*_ for transformation into comparable units; we use this latter nomenclature below.

The second virial coefficient *BM*_1_ is a measure of macromolecular interactions in solution and is proportional to the sum of the potential of mean force over all separations and orientations (47,48). This potential includes separation caused by excluded volume effects, specific attractive interactions such as electrostatic, hydrophobic, and hydrogen bonding, and electrostatic repulsive interactions. A positive *BM*_*1*_ indicates solute repulsion or large volume occupancy, a negative *BM*_*1*_ indicates solute self-association or volume compression. We find that for uncharged solutes like PEG, or PEG-HSA conjugates in high salt concentration, excluded volume is the dominant component of the second virial coefficient. Here we calculate the second virial coefficient due to excluded volume *BM*_*EX*_ as derived by Tanford (49),

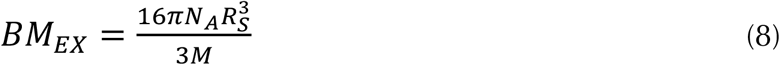

where the symbols are as defined as in Eq. 4.

As an additional approach to quantify experimentally determined solution non-ideality, sedimentation and diffusion coefficients can be fit to the following phenomenological equations as described in the companion paper (8),

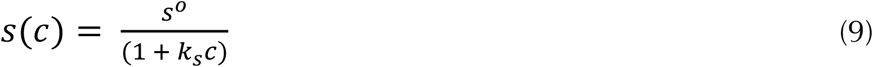

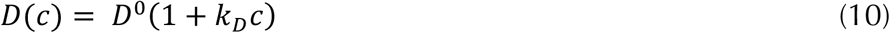

where *s(c)* is the measured sedimentation coefficient at concentration *c* in mg/ml, *s*^*0*^ is the sedimentation coefficient at infinite dilution, *k*_*S*_ is the hydrodynamic nonideality or change in sedimentation rate with concentration, *D(c)* is the diffusion coefficient measured by DLS at concentration *c, D*^*0*^ is the diffusion coefficient at infinite dilution, and *k*_*D*_ is the change in diffusion coefficient with concentration (50,51). The relationship of *BM*_EX_ to *k*_*S*_ and *k*_*D*_ is discussed below. The experimental determination of both *s*^*0*^ and *D*^*0*^ require performing multiple measurements with a range of concentrations and it would be advantageous to calculate these coefficients from structure, or as in this case, from structural ensembles.

### Calculation of Sedimentation Velocity Non-ideality from Hydration and Frictional Drag

Sedimentation velocity non-ideality *k*_*S*_ is determined by hydrodynamic backflow and frictional drag. Hydrodynamic backflow is the phenomena that as a particle sediments in solution, the solvent must flow counter to the sedimenting particle to fill in the vacated space. Both backflow and frictional drag are influenced by neighboring macromolecules.

The concentration dependence of sedimentation coefficients determined by sedimentation velocity has been calculated historically using the expression (Eq. 11) described by Rowe (34) that relates the hydrodynamic non-ideality constant *k*_*S*_ to the ratios 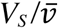 and *f/f*_*0*_.

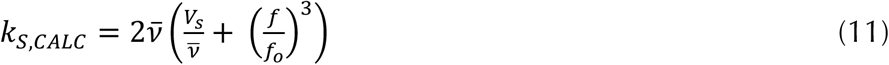

where *V*_*S*_ is the hydrated molecule specific volume (swollen volume), 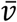 is the anhydrous partial specific volume and *f*/*f*_0_ is the frictional ratio of the hydrated, aspheric particle relative to a sphere of the same anhydrous molecular volume.

A calculated *k*_*S*_ using Eq. 11 was compared to experimentally determined *k*_*S*_ using Eq. 9 for the PEG, HSA and PEG-HSA species as reported in the companion paper (8). The data are plotted in Fig. 9 (grey circles). The calculated values are uniformly less than the experimental values indicating that the increased frictional drag on sedimentation with increased concentration was not completely accounted for in Eq. 11. To account for additional frictional drag we added a term for species specific intrinsic viscosity [*η*] as in Eq. 12. Note that this additional empirical term has the same units as *k*_*S*_ (ml/g).

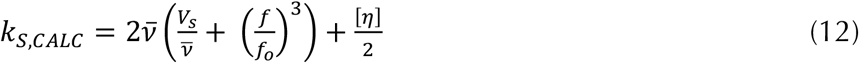

**Figure 9.**
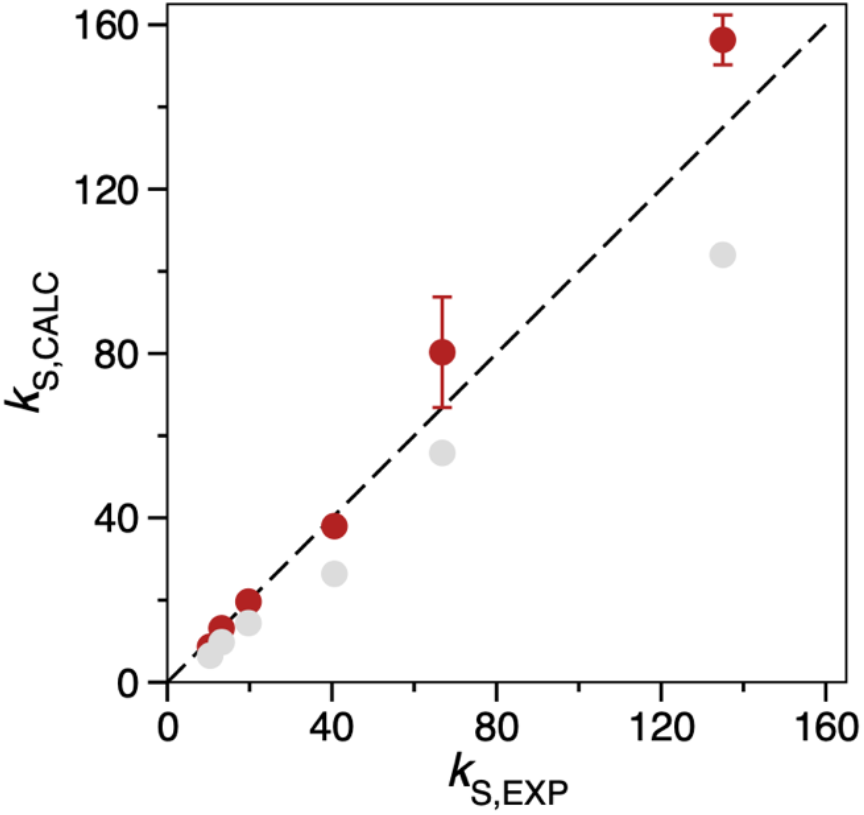
Comparison of methods to calculate sedimentation non-ideality constants. The data for ensemble calculated and experimental *k*_*S*_ values in Table S3 are plotted against each other as circles. Red, calculated by Eq. 12 (with viscosity correction); grey, calculated by Eq. 11 (original Rowe equation). Standard deviations are shown as capped error bars; some error bars are smaller than the data circle. The dashed line has a slope of 1.0 and intercept of zero.

Incorporating a term for intrinsic viscosity improves the agreement between experimentally determined and calculated *k*_*S*_ for HSA alone, and also improves the agreement with the experimental *k*_*S*_ values for the PEG and PEG-HSA conjugates (Fig. 9, red circles and Table S3, first three data columns). Dividing the [*η*] term by 2 optimized the agreement. The inability of Eq. 11 to predict experimental *k*_*S*_ values does not appear to be due to inaccuracies in the calculated values of 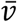, *V*_*S*_, or *f*/*f*_0_. The model calculated *f*/*f*_0_ values agree with those determined experimentally (Table S4). Both *V*_*S*_ and 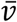 are core calculations in HullRad, the first from the convex hull volume and the second from widely accepted atomic volumes (52,53). These cannot be altered without causing errors in the overall calculation of hydrodynamic properties. The improvement in agreement between the calculated and experimental *k*_S_ values with Eq. 12 suggests that when calculating *k*_S_ an additional term for solute induced viscosity should be included.

### Calculation of Diffusion Non-ideality from Hydration and Frictional Drag

The concentration dependence of diffusion coefficients was calculated from model ensembles using a variation of the expression described by Teraoka (54),

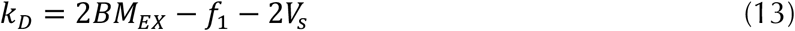

where *k*_*D*_ is the diffusion non-ideality constant, *BM*_*EX*_ is the excluded volume second virial coefficient defined above, *f* is the first order concentration frictional coefficient, and *V*_*S*_ is the hydrated molecule specific volume as described above.

The first order concentration frictional coefficient *f* is defined as,

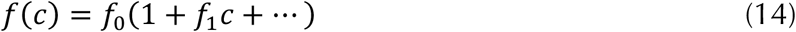

where *f(c)* is the frictional coefficient at concentration *c*, and *f*_*0*_ is the frictional coefficient at infinite dilution. In the absence of experimental frictional coefficient values in concentrated solutions we used the relationship *f*= [*η*] as an approximation (55).

The substitution here of a solute induced viscosity correction for frictional drag has precedent. Previous model calculations of the diffusion coefficient for particles in concentrated solutions have included corrections for apparent viscosity in different ways: The diffusion coefficient of particles in concentrated solutions may be accurately calculated from Eq. 3 by replacing the solvent viscosity *η* with the solution viscosity *η*_*φ*_ at volume fraction *φ* (56); alternately, a “hydrodynamic” correction for increased drag due to the flow induced by nearby particles has been added to the calculation (57). We use the intrinsic viscosity [*η*] as a proxy for increased friction in concentrated solutions.

Using Eq. 13 (with [*η*] as a substitution for *f*_*1*_), the calculated diffusion non-ideality constants *k*_*D*,*CALC*_ are in reasonable agreement with the experimental values *k*_*D*,*EXP*_ calculated from Eq. 10 (8) as shown in Fig. 10 (red circles) and Table S3.

**Figure 10.**
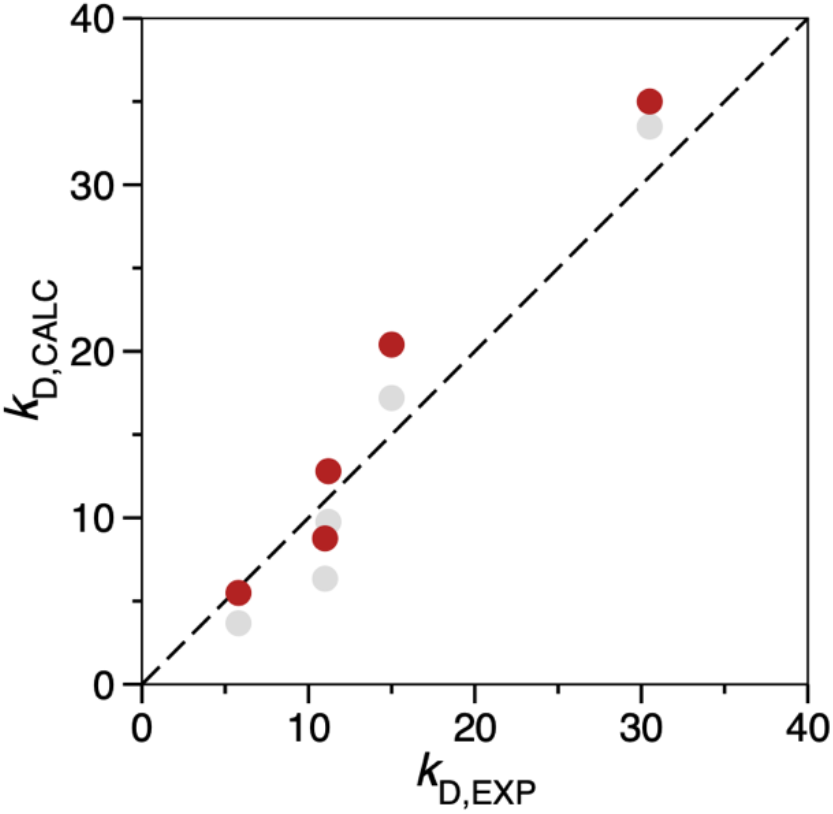
Ensemble calculated diffusion non-ideality constants vary with expression used. The data for ensemble calculated and experimental *k*_*D*_ values in Table S3 are plotted against each other. Red circles, Eq. 12 (modified); grey circles, Eq. 15 (modified). The dashed line has a slope of 1.0 and intercept of zero.

An alternative to Eq. 13 for the diffusion non-ideality constant was derived by Harding and Johnson (58),

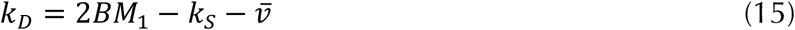

where, as above, *BM*_*1*_ is the sedimentation thermodynamic second virial coefficient, *k*_S_ is the sedimentation non-ideality constant, and 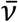 is the anhydrous partial specific volume. We tested Eq. 15 using *BM*_*EX*_ as a substitute for *BM*_*1*_ and *k*_S_ as defined by Eq. 12 (with intrinsic viscosity correction). The results are shown as grey circles in Fig. 10. Both Eqs. 13 and 15 provide reasonable estimates for the diffusion non-ideality constant for HSA and PEG-HSA.

### Estimating the Effect of Electrostatic Repulsion on the Second Virial Coefficient

The fact that *BM*_*EX*_ appears to be a good estimate of the second virial coefficient in Eqs. 13 and 15 indicates that excluded volume is the dominant effect of concentration on diffusion non-ideality. This result is surprising because the net charge on HSA in PBS has been estimated to be -16.3e (44) to -17.2e (Personal communication, Tom Laue). Although the experimental values for *k*_*D*,*EXP*_ were obtained in phosphate buffered saline where electrostatic repulsion would be significantly screened (30), some electrostatic repulsion would be expected in addition to the excluded volume particle separation. In this latter case the total calculated second virial would be the sum of both excluded volume (*BM*_*EX*_) and electrostatic repulsion (*BM*_*Z*_) terms, *BM*_*1*,*CALC*_ = *BM*_*EX*_ + *BM*_*Z*_.

Our estimate of *BM*_EX_ is consistent with available data. Table S5 lists the calculated values of 2*BM*_*EX*_ for HSA, PEG and PEG-HSA studied here. Sønderby et al. obtained a 2*BM*_EX_ of 14.1 mg/ml for recombinant HSA at pH 7 in high ionic strength buffer (∼600 mM NaCl) from fitting to static light scattering data (44). This is close to our values of 13.1 (Table S5) and 13.4 in the companion paper (8).

In contrast, accurate calculation of the second virial due to electrostatic interaction (*BM*_*Z*_) is difficult (59). Wills and Winzor (60) derived an expression from McMillan-Mayer theory (48) that has been used in the AUC community (61). An alternate expression from Tanford gives essentially the same values for the calculated electrostatic second virial (49),

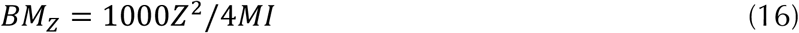

where *Z* is the net molecular charge, *M* is the molecular mass, and *I* is the ionic strength. The calculated 2*BM*_*Z*_ for HSA using Eq. 16 and the ionic strength of PBS is 13.7 ml/g. This value is significantly larger than the measured 2*BM*_*Z*_ found by Sønderby et al. (4.52 ml/g) at the equivalent salt concentration of PBS assuming an HSA net charge of -16.3e (44). It has been well documented that using the above expressions derived from theory may overestimate the extent of electrostatic repulsion (62,63). In fact, solvation forces may cause negative particles to be attractive at close approach (64). The 2*BM*_*Z*_ measured by Sønderby et al. (4.52 ml/g) is the expected value from Eq. 16 at an ionic strength equal to 0.41 M. Using *I*=0.41 in Eq. 16 we calculated the predicted 2*BM*_*Z*_ for each of the PEG-HSA species studied here and these are listed in Table S5. These calculated values of 2*BM*_*Z*_ are relatively small compared to the 2*BM*_*EX*_ values and the trend is for less electrostatic repulsion as the size of the PEG-HSA increases. This result is due to the mass in the denominator of Eq. 16 but would also be expected from partial screening of charge from PEG. In any case, inclusion of electrostatic repulsion in the second virial (i.e., 2*BM*_*1*,*CALC*_ = 2*BM*_*EX*_ + 2*BM*_*Z*_) in Eqs. 12 and 15 would increase the calculated *k*_*D*_ values, especially for the lower molecular weight species, and result in worse agreement with experimental values (cf. Fig. 10).

In summary, we are able to use expressions containing parameters (*V*_*S*_, *f/f*_0_, [*η*], *BM*_*EX*_) that depend only on molecular hydration volume and shape to accurately calculate solution non-ideality for PEGylated HSA.

### Test of the Harding Equation

One of the goals in the companion experimental paper was to test the validity of a relationship derived by Harding and Johnson that relates hydrodynamic and thermodynamic non-ideality (58). This expression is a rearrangement of Eq. 15 and specifically equates the thermodynamic second virial coefficient with the sum of sedimentation and diffusion non-ideality constants, 2*BM*_1_ = *k*_S_ + *k*_D_ (assuming 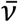 is much smaller than either *k*_S_ or *k*_D_).

We found that *BM*_EX_ is a good substitute for *BM*_1_ when calculating *k*_D_ for the molecules studied here (Fig. 10) and therefore tested the Harding expression using *BM*_EX_. The excluded volume second virial 2*BM*_*EX*_ shows excellent correlation with the ensemble calculated sums of *k*_S_ + *k*_D_, (Fig. S8, red circles, r = 0.999). Also included in Fig. S8 is the same plot with the sums of experimental *k*_S_ + *k*_D_ (8).

In the companion paper (8) the thermodynamic second virial coefficients determined by sedimentation analysis were significantly larger than the sums of *k*_S_ + *k*_D_ for all molecules tested and did not support the Harding equation. The good correlation between the sum of *k*_S_ + *k*_D_ and model calculated 2*BM*_*EX*_ illustrated in Fig. S8 does support the Harding equation when considering only excluded volume. These results suggest that other factors in addition to excluded volume may influence the thermodynamic second virial coefficient determined by sedimentation analysis.

## CONCLUSIONS

The sedimentation and diffusion properties of PEGylated HSA at infinite dilution are calculated from model ensembles with a high degree of accuracy when using the HullRad algorithm that takes into account hydration volume and shape. This agreement is shown in Fig. 4 above and emphasized in Fig. 8A in the companion paper (8). Only simple force field terms, together with fast coarse-grained simulations, are necessary to create the model ensembles and this enables analysis of large random coil polymers. Structural analysis with ensemble averaging appears to be a reasonable and sufficient approach to calculation of PEG random coil properties in a good solvent.

One of the methods used to validate protein therapeutics is the determination of sedimentation coefficients by analytical ultracentrifugation (65). The results presented here demonstrate that, even if the protein exhibits conformational variation, or is conjugated with a flexible polymer (66), it is possible to predict the measured sedimentation coefficient for aid in identifying molecular species and colligative properties that impact on formulation.

The solution concentration dependent properties of PEGylated HSA, as reflected in sedimentation and diffusion non-ideality constants, are a result of combined hydrodynamic interactions between a non-structured random coil polymer and a structured natively folded protein. In addition, there is an unexpected complex relationship between volume and shape, and hydrodynamic properties such as sedimentation coefficient and radius of gyration for these structurally heterogeneous molecules. We have developed expressions that accurately predict both fundamental hydrodynamic properties and concentration dependent non-ideality of PEGylated HSA in physiological salt from structure and these have been incorporated into HullRad.

## Authors contributions

P.J.F. and J.J.C designed the work; P.J.F. ran the simulations and calculated the results; P.J.F., J.J.C. and K.G.F. wrote the manuscript. All authors read and approved the final manuscript.

## Availability of data and materials

All calculated data are included in the text or supplementary information. Model ensembles are available upon request.

## Code availability

Code for HullRad is available at the HullRad website (hullrad.wordpress.com, hullrad.jhu.edu) and GitHub (github.com/fleming12/hullrad.github.io). CafeMol is available at the CafeMol website (www.cafemol.org).

## Acknowledgements

We would like to thank Drs. James Cole, J. Dave Dignam, Thomas Laue, Walter Stafford and Yaojun Zhang for helpful discussions and the Fleming lab members for critical feedback.

