## Supplemental Information for "The Molecular Basis for Hydrodynamic Properties of PEGylated Human Serum Albumin"

### SUPPLEMENTAL FIGURES

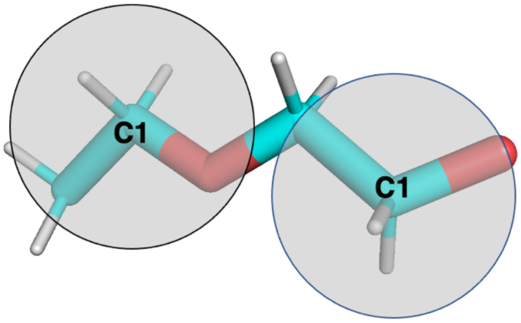


**Figure S1**. Polyethylene glycol coarse-grained model. Colored stick all-atom model of di-ethylene glycol with grey circles indicating positions of pseudo-atoms representing each ethylene oxide group. PyMOL (1) was used to create the image.

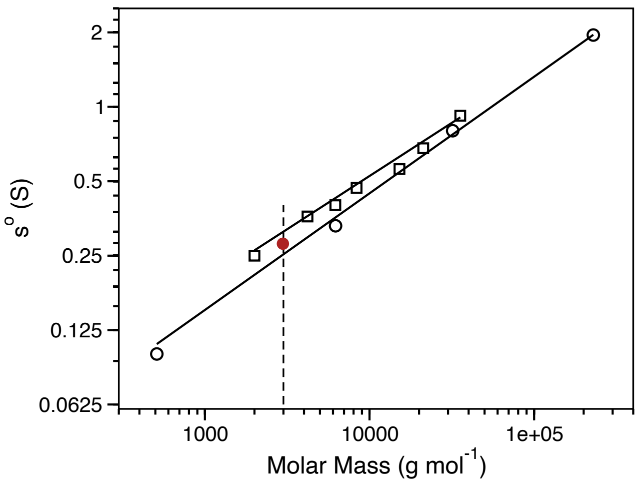


**Figure S2**. Experimental sedimentation coefficients were used to calibrate PEG model parameters. The plot shows corrected sedimentation coefficients ($s^o$) for different sized PEG from two published studies: Open square data from Nishchang et al. (2), open circle data from Luo et al. (3). The solid lines are least squares best fits to each data set. An average sedimentation coefficient of 0.281 (red circle) was used for parameterization of the coarse-grained simulation model of PEG68 (68 units, ~3000 g/mol)

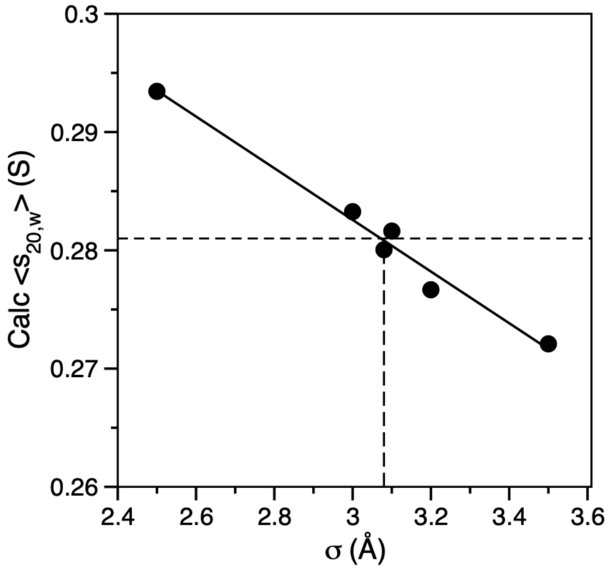


**Figure S3**. Calibration of the PEG coarse-grained excluded volume parameter. CafeMol simulations of PEG68 with various excluded volume parameters between 2.5 and 3.5 were run and the ensemble average sedimentation coefficients were calculated with HullRad (black circles). The solid black line is a least squares best fit to the data; the horizontal dashed line is the target sedimentation coefficient of 0.281; the vertical dashed line indicates that an excluded volume parameter of 3.08 generates a PEG ensemble that agrees with the average experimental sedimentation coefficient.

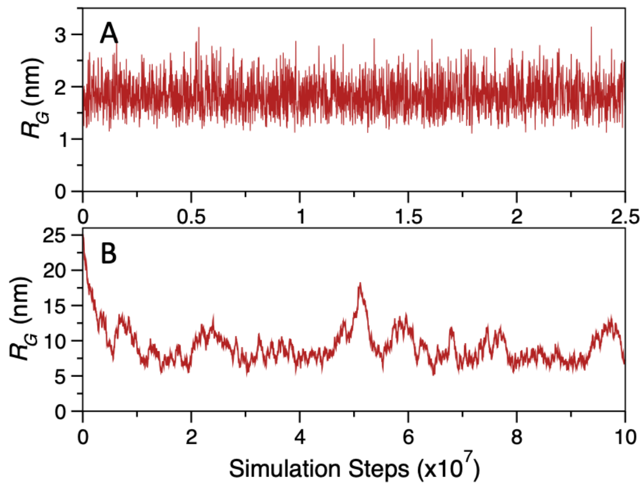


**Figure S4**. Time evolution of calculated $R_G$ for PEG68 and PEG908 simulations. (A) A coarse-grained model of PEG68 was simulated for 2.5 x$10^7$ steps. A thousand frames were evenly collected as PDB files and anhydrous radii of gyration calculated with HullRad (red line). The $R_G$ for the starting extended PEG68 structure is 7.1 nm and the structure is collapsed by the first sampling time. (B) A coarse-grained model of PEG908 was simulated for 1.0 x$10^8$ steps. A thousand frames were evenly collected and anhydrous radii of gyration calculated with HullRad (red line). Equilibrium is reached during the first 20 percent of the trajectory.

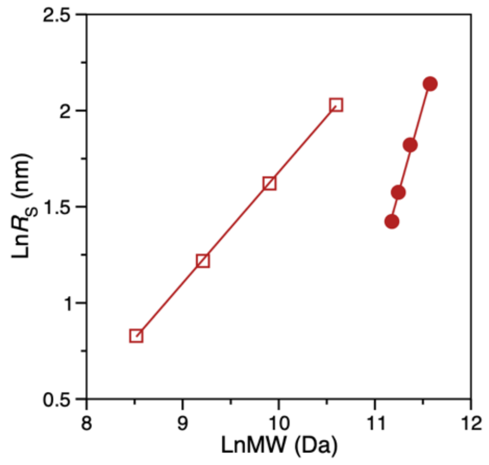


**Figure S5**. PEG-HSA conjugates do not follow pure random coil scaling laws. Log-log plot of Stokes radius $R_S$ against molecular weight. The open squares are for PEG alone; the solid circles are for the corresponding PEG-HSA conjugates. The solid lines are linear regressions with the slopes being equivalent to Flory scaling law exponents of 0.578 PEG alone (open squares), and 1.78 for PEG-HSA conjugates (solid circles).

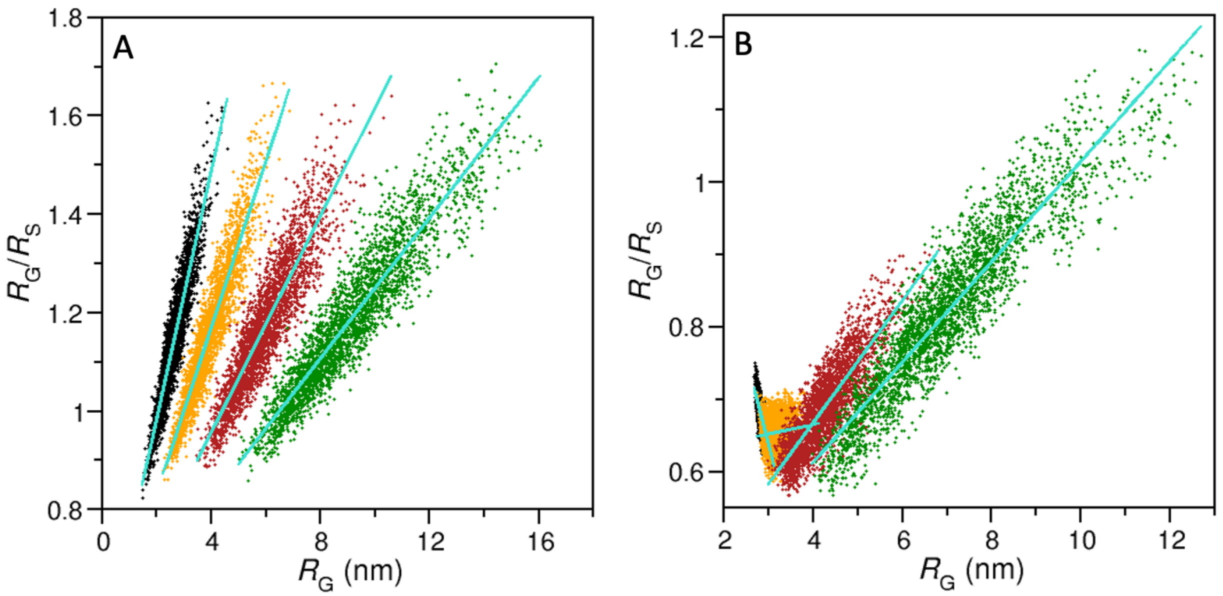


**Figure S6**. The $R_G/R_S$ ratio is size and structure related. The $R_G/R_S$ ratio is plotted against $R_G$ for (A) the PEG only moieties and (B) the complete PEG-HSA conjugates. Black circles, 5K PEG; orange, 10K PEG; red, 20K PEG; green, 40K PEG. The cyan solid lines are least squares fits to the data with slopes: (A, left to right); 0.252, 0.168, 0.110, 0.071; (B, left to right); -0.238, 0.013, 0.045,0.069.

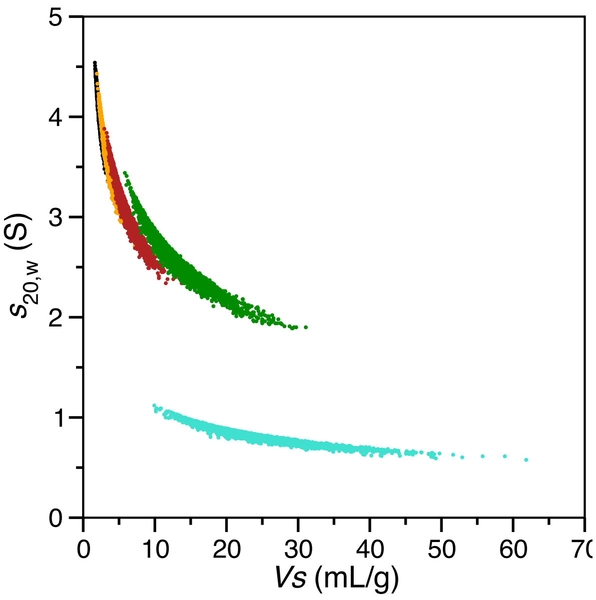


**Figure S7**. Non-linear dependence of sedimentation coefficient on hydrated volume. The individual model calculated $s_{20,W}$ values are plotted against the corresponding hydrated (swollen) volumes $V_S$ for combined ensembles (N=3000) of PEG and PEG-HSA conjugates as filled circles. Black, 5K PEG-HSA; orange, 10K PEG-HSA; red, 20K PEG-HSA; green, 40K PEG-HSA; cyan, 40K PEG alone.

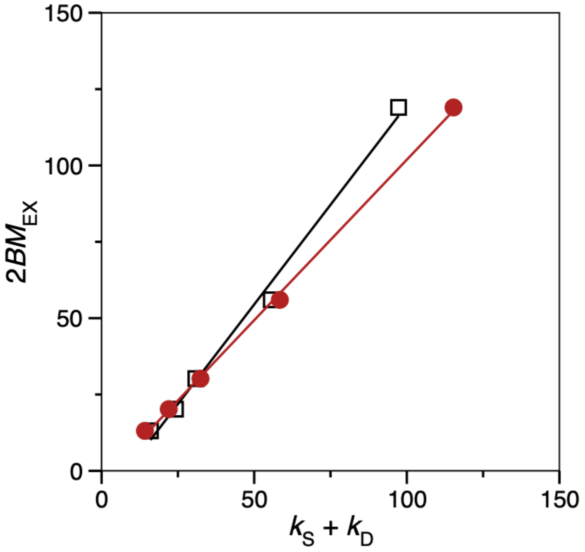


**Figure S8**. The sum of $k_S + k_D$ is correlated with excluded volume second virial coefficient. The ensemble calculated sums $k_S + k_D$ (red solid circles) and the corresponding experimentally determined sums from the companion paper (4) (black open squares) are plotted versus the excluded volume second virial coefficient ($2BM_{EX}$) calculated from model ensembles as listed in Table S5. Lines are linear regressions with (red circles) $r = 0.999$, slope = 1.05, intercept = -3.35; (black squares) $r = 0.997$, slope = 1.31, intercept = -10.8.

### SUPPLEMENTAL TABLES

**Table S1** Calculated ensemble average PEG radius of gyration agrees with experiment

| Sample | $R_G$ (nm) <Calc.>[a] | $R_G$ (nm) Exp.[b] | $R_G$ (nm) Exp.[c] |
|---|---|---|---|
| PEG77 (3400 MW) | 1.97 (0.05) | 2.10 | 1.91 |

[a]Mean of three ensembles, standard deviation in parentheses.
[b]Neutron scattering, (5)
[c]Neutron scattering, (6)

**Table S2** Calculated PEG and PEG-HSA hydrodynamic properties agree with experimental values

| Sample | $\langle R_S \rangle$ (nm) Exp.[a] | $\langle R_S \rangle$ (nm) Calc.[b] | $\langle s_{20,W} \rangle$ (S) Exp.[c] | $\langle s_{20,W} \rangle$ (S) Calc.[b] | $D_{20,W}$ ($cm^2\ sec^{-1}$) Exp.[d] | $\langle D_{20,W} \rangle$ ($cm^2\ sec^{-1}$) Calc.[b] |
|---|---|---|---|---|---|---|
| 40K PEG | 7.48 | 7.53 (0.08) | 0.81 | 0.82 (0.01) | ND | 2.87 |
| 40K PEG-HSA | 8.17 (0.24) | 8.49 (0.39) | 2.70 | 2.59 (0.11) | 2.54 | 2.55 (0.11) |
| 20K PEG-HSA | 5.97 (0.06) | 6.18 (0.02) | 3.15 | 3.05 (0.01) | 3.55 | 3.49 (0.01) |
| 10K PEG-HSA | 4.83 (0.04) | 4.83 (0.04) | 3.53 | 3.57 (0.03) | 4.48 | 4.46 (0.04 |
| 5K PEG-HSA | 4.14 (0.01) | 4.15 (0.01) | 3.97 | 3.97 (0.01) | 5.16 | 5.18 (0.01) |

[a]Mean of experimental values obtained by sedimentation (4). Numbers in parentheses are standard deviations for three independent experiments.
[b]Mean of model ensembles. Numbers in parentheses are standard deviations for three independent ensembles.
[c]Mean of two experimental values obtained by sedimentation (4).
[d]Determined by DLS (4)

**Table S3** HSA and PEG-HSA concentration dependent coefficients calculated with alternative expressions

| Sample | $k_S$ **Exp.**[a] | $k_S$_Fl Calc.[b] | $k_S$_Rw Calc.[c] | $k_D$ **Exp.**[a] | $k_D$_A Calc.[d] | $k_D$_B Calc.[e] |
|---|---|---|---|---|---|---|
| 40K PEG | **135** | 156 | 104 | **-** | - | - |
| 40K PEG-HSA | **66.8** | 80.3 | 55.8 | **30.5** | 35.0 | 33.5 |
| 20K PEG-HSA | **40.6** | 38.0 | 26.4 | **15.0** | 20.4 | 17.2 |
| 10K PEG-HSA | **19.7** | 19.6 | 14.4 | **11.2** | 12.8 | 9.76 |
| 5K PEG-HSA | **13.2** | 13.2 | 9.76 | **11.0** | 8.76 | 6.34 |
| HSA | **10.4** | 9.64 | 6.48 | **5.8** | 5.49 | 3.67 |

[a]Mean of experimental values obtained by sedimentation (4).
[b]ks_Fl = $2\bar{v}\left(\frac{V_s}{\bar{v}}+\left(\frac{f}{f_o}\right)^3\right)+\frac{[\eta]}{2}$ (Eq. 11, Add intrinsic viscosity term)
[c]ks_Rw = $2\bar{v}\left(\frac{V_s}{\bar{v}}+\left(\frac{f}{f_o}\right)^3\right)$ (Eq. 10)
[d]kd_A = $2BM_{EX}-[\eta]-2V_s$ (Eq. 12, Modified)
[e]kd_B = $2BM_{EX}-k_{S\_Fl}-\bar{v}$ (Eq. 16, Modified)
All calculated values are from combined ensembles, *N*=3000. Equation numbers refer to the main text.

**Table S4** Comparison of methods to calculate frictional ratios

| Sample | $f/f_0$ Exp[a] | $<f/f_0>$ Calc.[b] |
|---|---|---|
| 40K PEG | 3.21 | 3.19 |
| 40K PEG-HSA | 2.51 | 2.66 |
| 20K PEG-HSA | 2.00 | 2.09 |
| 10K PEG-HSA | 1.71 | 1.71 |
| 5K PEG-HSA | 1.49 | 1.50 |
| HSA | 1.31 | 1.31 |

[a]Calculated as described in companion paper (4).
[b]Mean of combined model ensembles calculated with HullRad.

**Table S5** Ensemble calculated second virial coefficients

| Sample | $2BM_{EX}$[a] Calc. (ml/g) | $2BM_{Z}$[b] Calc. (ml/g) | $2BM_{1,CALC}$[c] Calc. (ml/g) |
|---|---|---|---|
| 40K PEG | 222 | 0.0 | 222 |
| 40K PEG-HSA | 119 | 1.69 | 121 |
| 20K PEG-HSA | 56.0 | 2.08 | 58.1 |
| 10K PEG-HSA | 30.2 | 2.36 | 32.6 |
| 5K PEG-HSA | 20.2 | 2.52 | 22.7 |
| HSA | 13.1 | 4.38 | 18.5 |

[a]Mean of combined model ensembles calculated with Eq. 13.
[b]Calculated with Eq. 16, $I$=0.41.
[c]From $2BM_{1,CALC} = 2BM_{EX} + 2BM_{Z}$.
Equation numbers refer to the main text.
